# Amino Acid Requirements of the Chinese Hamster Ovary Cell Metabolism during Recombinant Protein Production

**DOI:** 10.1101/796490

**Authors:** Bergthor Traustason

## Abstract

Majority of biopharmaceutical drugs today are produced by Chinese hamster ovary (CHO) cells, which have been the standard industry host for the past decades. To produce and secrete a substantial amount of the target recombinant proteins the CHO cells must be provided with suitable growth conditions and provided with the necessary nutrients. Amino acids play a key role in this as the building blocks of proteins, playing important roles in a large number of metabolic pathways and being important sources of nitrogen as well as carbon under certain conditions. In this study exploratory analysis of the amino acid requirements of CHO cells was carried out using metabolic modelling approaches. Flux balance analysis was employed to evaluate the optimal distribution of fluxes in a genome-scale model of CHO cells to gain information on the cells’ metabolic response *in silico*.

The results showed that providing non-essential amino acids (NEAAs) has a positive effect on CHO cell biomass production and that cysteine as well as tyrosine play a fundamental role in this. This implies that extracellular provision of NEAAs limits the extent of energy loss in amino acid biosynthetic pathways and renders additional reducing power available for other biological processes. Detailed analysis of the possible secretion and uptake of D-serine in the CHO model was also performed and its influence on the rest of the metabolism mapped out, which revealed results matching various existing literature. This is interesting since no mention of D-serine in regard to CHO cells was found in current literature, as well as the fact that this opens up the possibility of using the model for better understanding of certain disorders in higher up organisms that have been implicated with D-serine, such as motor neuron and cognitive degeneration. Finally, outcome from the model optimisation of different recombinant proteins demonstrated clearly how the difference in protein structure and size can influence the production outcome. These results show that systematic and model-based approaches have great potential for broad *de novo* exploration as well as being able to handle the cellular burden associated with the production of different types of recombinant protein.

## 1 Introduction

The biopharmaceutical market is growing rapidly worldwide with the emergence of biopharmaceuticals as widely used treatments for numerous diseases [1]. The global biopharmaceuticals market was valued at $162 billion in 2014 and is expected to reach $278 billion by 2020 [2, 3]. Majority of biopharmaceutical drugs are produced by Chinese hamster ovary (CHO) cells, which have remained the standard industry host for the past three decades [4]. CHO cells are predominantly used as expression hosts for recombinant monoclonal antibody (mAb) production [1, 5], which is also the fastest growing segment of the biopharmaceutical industry [6]. The reason for this wide use of CHO cells is mainly their ability to perform post-translational modifications and correctly fold recombinant proteins, as well as their proven track-record of being safe hosts [4, 5].

To produce and secrete a substantial amount of the target recombinant protein the CHO cells must be provided with suitable growth conditions. The nutrients provided in the culture medium is one of the most important factors in establishing this optimal growth environment for the cells and a fundamental part in the design of upstream cell culture biopharmaceutical processes [7]. Culture medium is usually made up of various carbon sources, lipids, minerals, vitamins, amino acids, hormones, growth factors and various other proteins [8, 9]. Amino acids constitute the building blocks of both recombinant and native protein, which make up *ca.* 70% of the cell’s dry mass [7, 10]. Mammalian cells can only produce some of the required amino acids for protein synthesis, nonessential amino acids (NEAAs), while others need to be provided externally, essential amino acids (EAAs). Amino acids act as precursors for a multitude of intermediates that play a role in a large number of metabolic pathways, as well as acting as important sources of nitrogen, and potentially as a source of carbon when other sources become limiting. Therefore, it is important that the cells are provided with the right amount of EAAs to survive as well as NEAAs to facilitate the favourable use of resources. This significance of amino acids in medium design has been recognised for a long time, however, even after decades of industrial practice, optimisation of the provided amino acids continues to have possibilities for improvement [11]. In fact, no official systematic procedure exists for CHO cell culture medium optimisation, and several strategies followed by the industry have led to the emergence of a vast number of different medium formulations predominantly developed based on relevant experience [12].

Two common experimental strategies used for this purpose are statistical design, which allows for statistical analysis and control of external variables, and metabolic profiling, which provides information on the physiology of the cell. These types of experimental methods can often be resource-intensive and require large amount of experiments [6]. Therefore, there is emerging interest in using modelling approaches to assist and guide the laboratory experiments to achieve further improvements. Most of these modelling approaches can roughly be defined as either kinetic or stoichiometric. Kinetic models establish relationships between metabolites through kinetic laws and are represented by differential-algebraic equations. Understanding the dynamic behaviour of the system is extremely useful in bio-process design and improvement. However, the disadvantage of kinetic models is that there are often complex non-linear equations that can render the solution process difficult. The relevant kinetic parameters are also often lacking an accurate value since it is generally difficult to recreate exactly the dynamic interactions among the various components in an intracellular environment [13].

### 1.1 Stoichiometric models

Stoichiometric models are one of the approaches used for the problem at hand. In principle they involve writing down all the relevant reactions that occur in a particular system and describing them mathematically. These reactions are identified and each of their fluxes is determined using methods such as metabolic flux analysis (MFA) and flux balance analysis (FBA) [14]. Both FBA and MFA constrain the network of reactions according to factors such as conservation of mass and energy, as well as observed conditions in the studied environment. These methods both rely on the assumption of pseudo-steady state, i.e. that accumulation of intracellular metabolites is negligible compared to the observed fluxes [14]. Usually, MFA is used with simpler networks that are fully defined by measured extracellular reactions. FBA, on the other hand, utilises all known reactions of the studied cells that can be found in literature sources and databases based on genomic data.

Xing *et al.* used MFA to optimise the amino acid composition of CHO cell culture media for manufacturing a mAb [15]. The MFA was performed using 32 specific metabolic rates derived from two semi-steady states of continuous culture. Modifications were formulated for the media based on both specific metabolic rates and fluxes. This resulted in modified media, enhancing peak cell density and the protein concentration without significantly impacting the quality. Various other reports use MFA similarly for CHO cells but do not focus on amino acids [16, 17].

Selvarasu *et al.* combined metabolomics and *in silico* modelling in an approach to gain new understanding of the intracellular mechanisms of CHO cultures [10]. A shortlist of metabolites associated with growth limitation was created based on metabolite profiling. A model representing the key metabolic functions of CHO cells was developed, which enabled characterisation of internal metabolic behaviours. It is good to keep in mind that stoichiometric models are condition specific, however, one of their main advantages is that they can account for competing reactions in the metabolic network. These methods also only work properly if the system reaches pseudo-steady state, this assumption is not a problem for industrial CHO cell culture since the process involves a time scale of days.

### 1.2 Flux Balance Analysis (FBA)

FBA is a mathematical approach to analyse flow of metabolites through a metabolic network [14]. Modelling frameworks can be represented by a stoichiometric matrix (**S**), where the rows represent the metabolites of the model and the columns the reactions, and a vector of reaction fluxes (**v**), indicating the reaction rates. Multiplying the stoichiometric matrix with the flux vector defines the mass balance for each of the metabolites of the model. By assuming pseudo-steady state all the mass balances can be set to zero and form a system of linear equations. Additional constraints can then be introduced by restricting fluxes with specific upper and lower bounds. An objective function (Z) is then defined, for example maximisation of biomass to analyse the growth of the cell. Linear programming can then be employed to find values for the fluxes in the model that fulfil the optimisation problem. This linear programming problem can be put forward mathematically as:

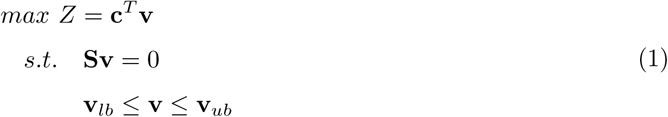

where **c** is a vector of the linear objective coefficients, **v**_*lb*_ are the set lower bounds and **v**_*ub*_ the set upper bounds. These upper and lower bounds along with the constraints **Sv** = 0, determine the feasible region of the problem.

### 1.3 Flux Variability Analysis (FVA)

The solution to an FBA problem is generally not a unique vector of optimal fluxes. That is, although solving the problem put forward in (1) returns an optimal flux vector, **v**^*∗*^ ∈ ℝ^*n*^, with a single value for each reaction, there is usually an infinite set of steady-state flux vectors that can satisfy the constraints while giving the same optimal results: **c**^*T*^**v**^*∗*^ = **c**^*T*^ **v**. Flux variability analysis (FVA) is a method to find the maximum and minimum range of each of the reaction fluxes that can still satisfy all the constraints [18]. This is performed using two optimisation problems for each reaction of interest, *v*_*i*_:

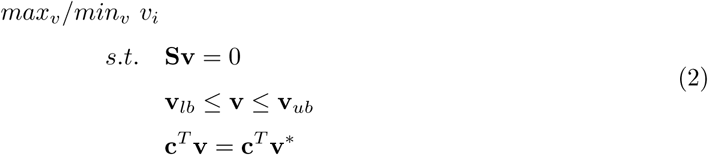

With this method it becomes possible to evaluate the robustness of metabolic models in varied simulation conditions.

Recently, Hefzi *et al.* reconstructed a consensus genome-scale metabolic network of a CHO cell building upon previous work carried out by other groups such as Selvarasu *et al.* The model, *iCHO1766*, has 1766 genes and 6663 reactions describing metabolism and protein production in CHO cells. It can, therefore, provide a platform for managing and interpreting CHO-relevant big data. Furthermore, with the enumeration of enzymes underlying metabolic pathways the reconstruction provides a mechanistic link between the phenotype and genotype. This allows for the effective integration of orthogonal data types, such as metabolomics, genetic variants, transcriptomics and growth rates. Along with the general model, cell-line specific models were also developed based on existing data used as input information in the GIMME algorithm [19]. These models are provided for CHO-S, CHO-K1, and CHO-DG44 cells. The models will be maintained and improved over time and allow for further studies on biotherapeutic production.

Calmels *et al.* manually adapted and tailored the model by Hefzi *et al.* to fit the metabolic profiles of high-yielding CHO cell lines used for industrial protein production as well as performing generic alterations to simplify the model and deal with missing metabolic constraints [20]. They silenced 537 transporters, greatly reducing the complexity of the model. This led to no more inconsistencies of solutions containing high flux loops that feed into each other being observed and led to disappearance of infeasible cycles linked to amino acid uptake fluxes [20]. They then constrained this updated model with 24 metabolites that were measured on a daily basis in four independent 2 L fed batch cell culture processes for industrial mAb production. With the model they could accurately predict both exometabolomic data and the growth rate. This shows how genome-scale models can provide insight into metabolism of cells cultivated in industrial processes.

In this study exploratory analysis of the amino acid requirement of CHO cells in the growth and production medium was carried out using metabolic modelling approaches. FBA was employed to evaluate the optimal distribution of fluxes in a genome-scale model of CHO cells to gain information on the cells’ metabolic response *in silico*. The metabolism of D-serine within the model was mapped out and its influence on the rest of the cell metabolism tested. This was then all put in perspective of recombinant protein production using CHO cells. Five different recombinant proteins were tested for production, four mAbs and one cyclotoxin fusion protein. This was then used to gain insight into how the media demand can vary depending on the different proteins being produced and help increase the understanding of the cellular metabolism.

## 2 Materials and Methods

In this study, analysis was carried out using metabolic modelling approaches. This was performed using COBRA, a software suite for quantitative prediction of cellular metabolic networks utilising constraint-based modelling [18]. FBA was employed on the genome-scale metabolic model of CHO cells developed by Hefzi *et al.* [19] and used to evaluate the metabolic response of the cell to variations in the extracellular availability of the medium, with a special emphasis on amino acids. FVA was then used to evaluate the range of calculated fluxes from FBA. The linear programming to perform both FBA and FVA was done in MATLAB 2018b [21], using the Gurobi optimiser [22].

### 2.1 Recombinant Proteins

Five different recombinant proteins were tested for production, four mAbs: alemtuzumab, blinatumomab, ibritumomab tieuxetan and trastuzumab, and one cyclotoxin fusion protein, denileukin diftitox. These proteins vary greatly in structure, size and functionality. They were chosen since information on them is readily available and they have been utilised and produced within industry. Some general technical information on each of the proteins can be found in Table 1, while the amino acid content for all the proteins can be found in Table 2. Further detailed information on each protein, such as on sequences and pharmacology, can be found on the DrugBank database [23].

**Table 1:**
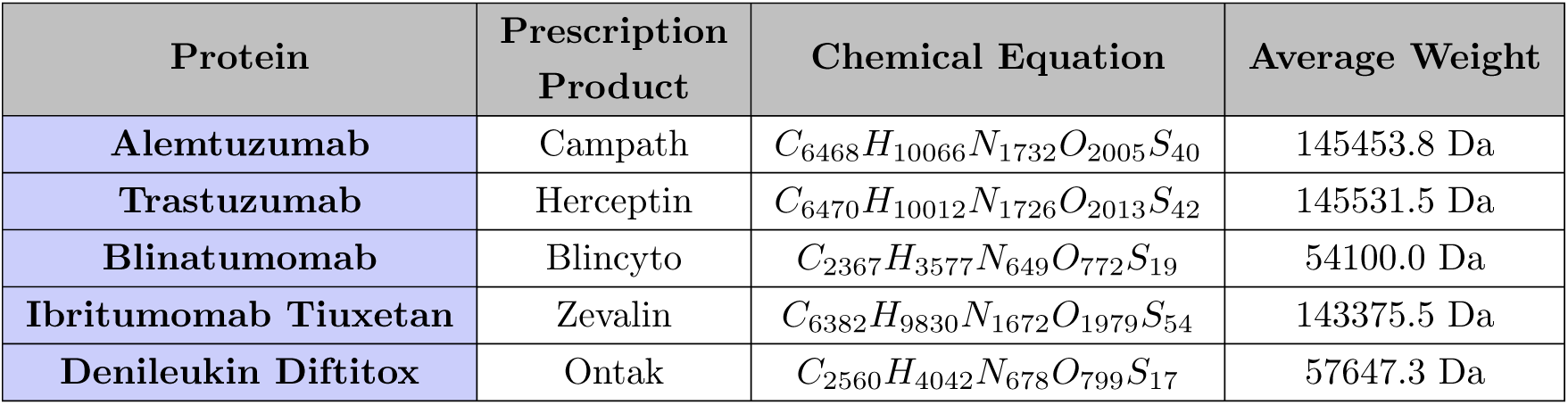
General technical information on the different recombinant proteins

**Table 2:** Number of amino acids for one molecule of each recombinant protein

| Amino Acid | Alemtuzumab | Trastuzumab | Blinatumomab | Ibritumomab<br>Tiuxetan | Denileukin<br>Diftitox |
| --- | --- | --- | --- | --- | --- |
| Alanine | 62 | 74 | 32 | 74 | 39 |
| Cystine | 30 | 32 | 9 | 32 | 5 |
| Aspartate | 56 | 56 | 24 | 52 | 23 |
| Glutamate | 58 | 62 | 16 | 60 | 43 |
| Phenylalanine | 48 | 44 | 10 | 40 | 18 |
| Glycine | 80 | 88 | 66 | 70 | 33 |
| Histidine | 26 | 26 | 10 | 26 | 12 |
| Isoleucine | 32 | 30 | 17 | 38 | 29 |
| Lysine | 94 | 90 | 25 | 84 | 39 |
| Leucine | 99 | 94 | 28 | 82 | 49 |
| Methionine | 10 | 12 | 10 | 22 | 12 |
| Asparagine | 56 | 50 | 11 | 52 | 31 |
| Proline | 96 | 92 | 19 | 94 | 19 |
| Glutamine | 66 | 62 | 31 | 56 | 19 |
| Arginine | 40 | 40 | 15 | 32 | 13 |
| Serine | 160 | 162 | 71 | 170 | 43 |
| Threonine | 116 | 110 | 41 | 110 | 34 |
| Valine | 120 | 120 | 23 | 120 | 37 |
| Tryptophan | 20 | 22 | 13 | 28 | 5 |
| Tyrosine | 54 | 62 | 33 | 62 | 18 |

### 2.2 Genome-Scale Model of CHO Cells

The genome-scale metabolic model developed by Hefzi *et al.* with the alterations adopted by Calmels *et al.* was used for the analysis carried out in this study. The necessary reaction pathways leading to the productions of the five recombinant proteins were added to the model. The structure and sequence of the five recombinant proteins was retrieved from the DrugBank database [23]. The energy requirements for the addition of an amino acid to an extended peptide chain in the assembly step were adapted from [24]. A general outline of this process can be seen in Figure 1.

**Figure 1:**
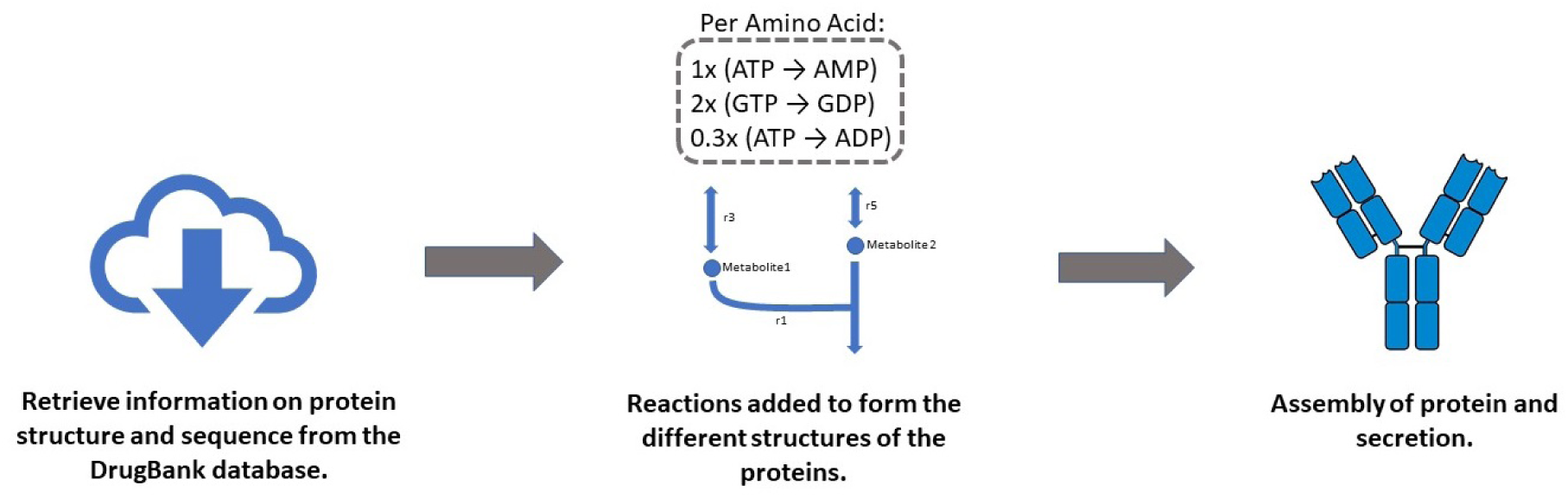
General outline of adding in the necessary reaction pathways leading to the production of the recombinant proteins.

A complete list of the modifications done according to Calmels *et al.* as well as details on each of the added recombinant protein pathways can be found in the supplementary material. A detailed comparison was also performed on all interconversions of amino acids in the model with the ones found in literature.

The model contains two biomass functions, one for a recombinant-protein producing cell line (*biomass_cho_producing*) and one for a nonproducing cell line (*biomass_cho*). The configurations for the functions were obtained based on literature values and both biomass reactions used experimentally determined amino acid composition [10]. The biomass function for the recombinant-protein producing cell line was used for all cases of biomass optimisation in this report.

### 2.3 Simulations and Model Setup

At first, various simulations were performed with most reactions of the model unbound while one or two uptake reactions were restricted at a time to get a general sense of their effect and the functionality of the model. After this, all specific cases for the model were done by restraining uptake reactions according to specific media. The benchmark constraints used for the simulations carried out were adapted from [19]. It is a serum-free medium which was used both with and without sodium butyrate (NaBu), which is often added to media to induce protein production. The one containing NaBu was selected since it showed the highest growth out of all the constraints that did not utilise temperature shifts and showed the smallest difference in model predictions and results from experiments [19].

To increase the robustness of the solutions from FBA, the model was further constrained. The model was reaching the maximum upper bound for the secretion of H_2_O and CO_2_. This meant that bicarbonate uptake was also reaching this maximum upper bound, since the organism tries to maintain balance in the pH levels through the bicarbonate-CO_2_ buffering system. In the literature, ranges for oxygen uptake and carbon dioxide production rates specifically for CHO cells are available [25]. The uptake rate of O_2_ given by the model using the benchmark medium constraints were, therefore, used to calculate what the corresponding range for CO_2_ secretion rates should be according to the literature. This was then used to further constrain the model, where the highest calculated CO_2_ secretion rate was set as the upper limit of the model reaction. All the applied constraints to simulate limits on nutrient availability in different media can be seen in the supplementary material.

An analysis of all the amino acids in the model and the effect it had on the biomass production to remove them was performed. The model was constrained according to the updated benchmark medium defined above like the rest of the simulations that can be found in this report, unless otherwise specified. The maximisation of biomass production was the objective function and each time an amino acid was removed the constraints were adjusted to replace its contents. All amino acids that made the biomass production decrease to zero were classified as EAA. A comparison was then made between the biomass production when the model was provided with all the amino acids and when it was only provided with the EAAs but with constraints configured to replace the contents of the removed NEAAs.

During the general unbound analysis, the solutions showed that the cell was secreting D-serine under certain extreme circumstances. This was considered very interesting since there is no previous mention of D-serine in relation to CHO cells in the literature. Further analysis was performed, first to see if the D-serine secretion would be observed under regular medium constrained conditions while various constraints were changed. Then the medium constrained cell was forced to secrete various amounts of D-serine while optimised for biomass production and these cases compared in an attempt to understand the metabolic change that takes place in the cell. FVA was used to determine if a detected flux difference between the cases was a notable change by looking at whether the maximum and minimum ranges of the reaction fluxes overlapped.

Finally, the model was optimised for the secretion of the five different recombinant proteins. This was done using the defined medium and no other restrictions as well as forcing the cell to produce at least a 10% of the maximum biomass production. This was also performed for the three different strain specific models. An alternative case was also tested where L-serine was replaced by D-serine in the defined medium and the model then optimised for recombinant protein production.

## 3 Results

### 3.1 Effect of NEAAs

According to the simulations there were nine amino acids essential in the model and eleven non-essential (Figure 2a). Removing all the NEAAs while replacing their contents in the medium resulted in a 41% decrease in the biomass production compared to the case containing all the amino acids (Figure 2b).

**Figure 2:**
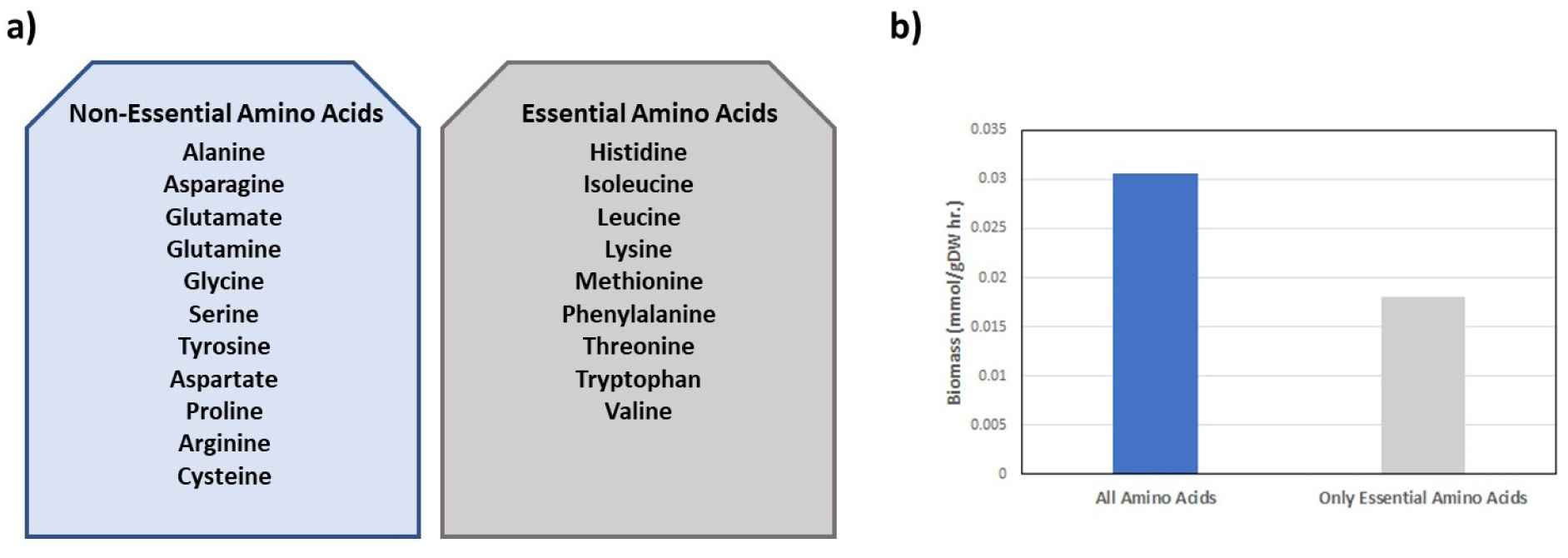
**a)** The NEAAs and EAAs in the general model. Tested by taking out each amino acid and replacing its content within the medium and determining for which the growth of the cell decreased to zero. **b)** The biomass production (growth), shown as mmol/gDW hr., for the defined medium constrained model containing all the amino acids or including only the EAAs, where the content of the removed NEAAs is still provided in the medium.

When removing the NEAAs one by one to test their effect on biomass, the removal of only two NEAAs had any effect on the amount of biomass produced. These amino acids were tyrosine and cysteine. The removal of all the other NEAAs had no notable effect on the biomass production (Figure 3).

**Figure 3:**
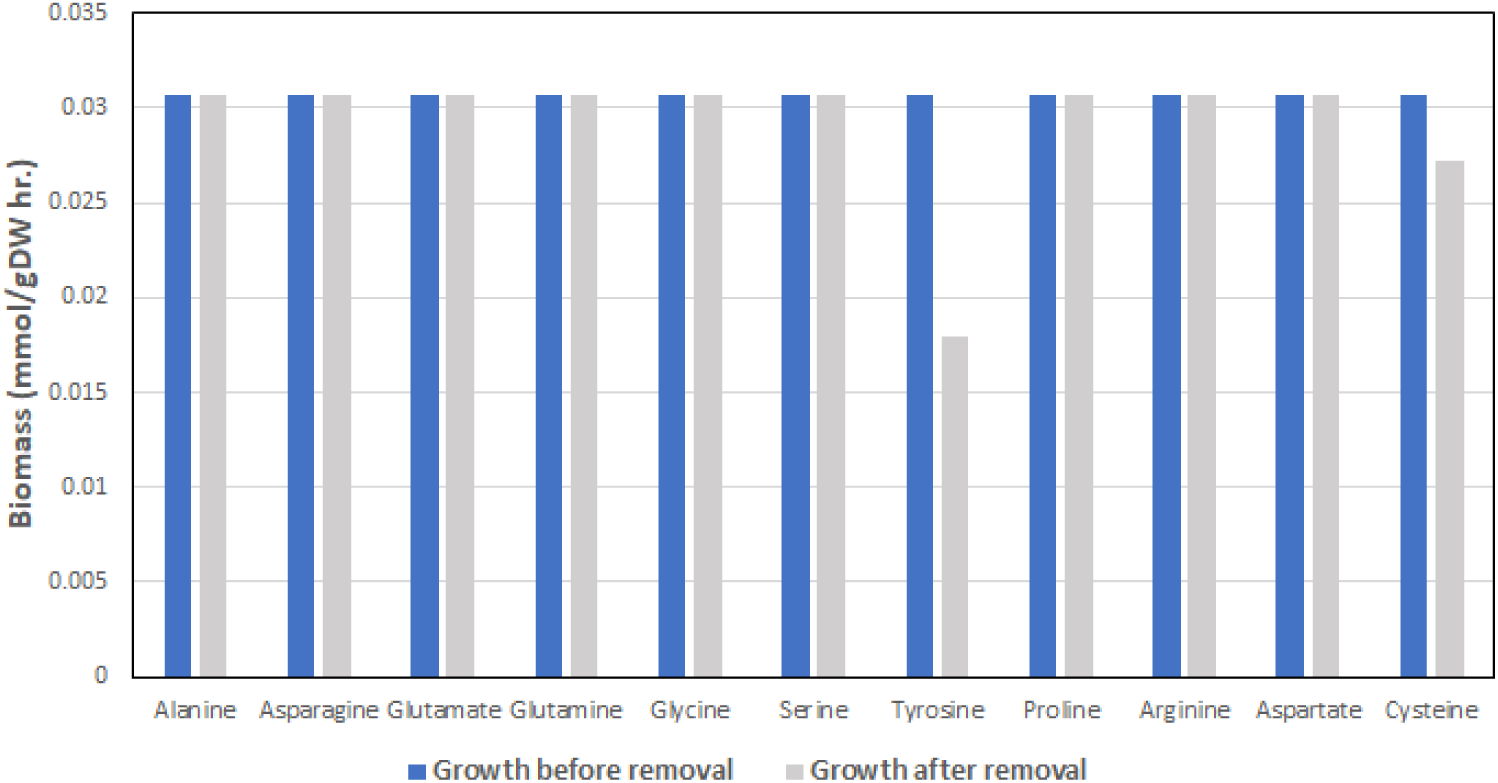
The effect on biomass production when uptake for each of the NEAAs is put to zero, while replacing their contents in the medium.

### 3.2 D-serine Secretion

After D-serine secretion was observed in some unbound analysis cases, different metabolic configurations were explored to determine whether the cell was able to secrete D-serine under defined medium constrained conditions. The uptake of glucose, glutamate, glutamine, glycine, L-serine, pyruvate and the secretion of CO_2_ were varied one by one as well as in combinations, since they were suspected to have the largest impact on the pathway leading to D-serine secretion. The fluxes of these reactions were varied ranging from 0 to the maximum model flux value, however, no D-serine secretion was detected in any of the cases.

The medium constrained cell was then forced to secrete various amounts of D-serine ranging from 0 to 0.382 mmol/gDW hr., which was the highest amount the cell could secrete before the optimisation problem became infeasible. The biomass production decreased as the D-serine secretion was increased (Figure 4). Excluding these two reactions, there were in total 30 reactions that showed a notable difference in fluxes between the secreting and non-secreting D-serine cases. Of these, 17 showed an increased flux while 13 decreased, as can be seen in Figures 5 and 6. The rest of the reactions did not show a notable difference in fluxes between the cases since their flux boundaries according to FVA overlapped. The net ATP consumption for the two cases was also compared, where the D-serine secreting case showed lower consumption.

**Figure 4:**
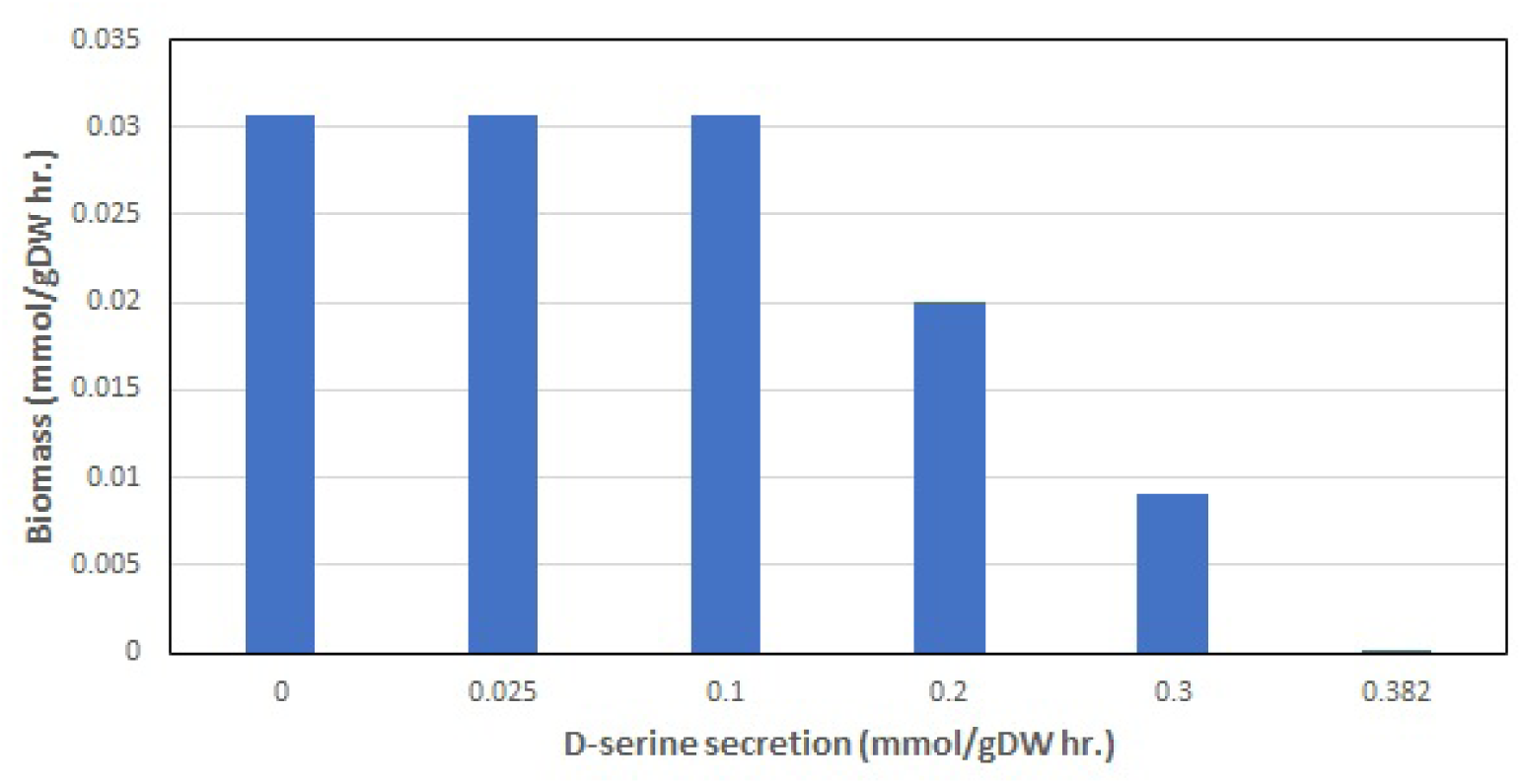
The effect on biomass production when D-serine secretion was increased.

**Figure 5:**
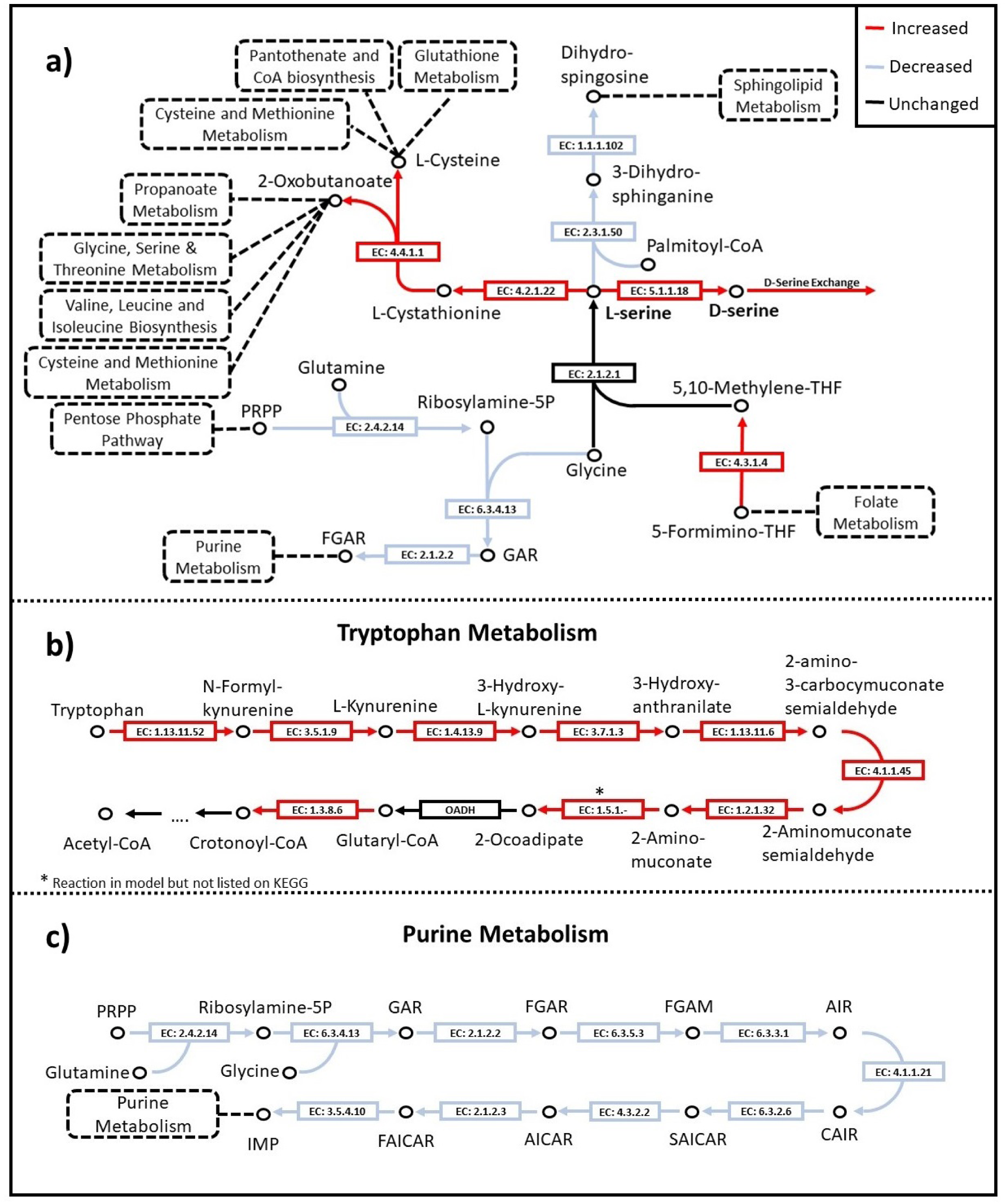
Reactions that show a notable flux change when the cell starts secreting D-serine. Reactions shown for: **a)** Pathways leading to L- and D-serine, **b)** Tryptophan metabolism, **c)** Purine metabolism. The two cases compared were when D-serine secretion was 0 mmol/gDW hr. and 0.2 mmol/gDW hr.

**Figure 6:**
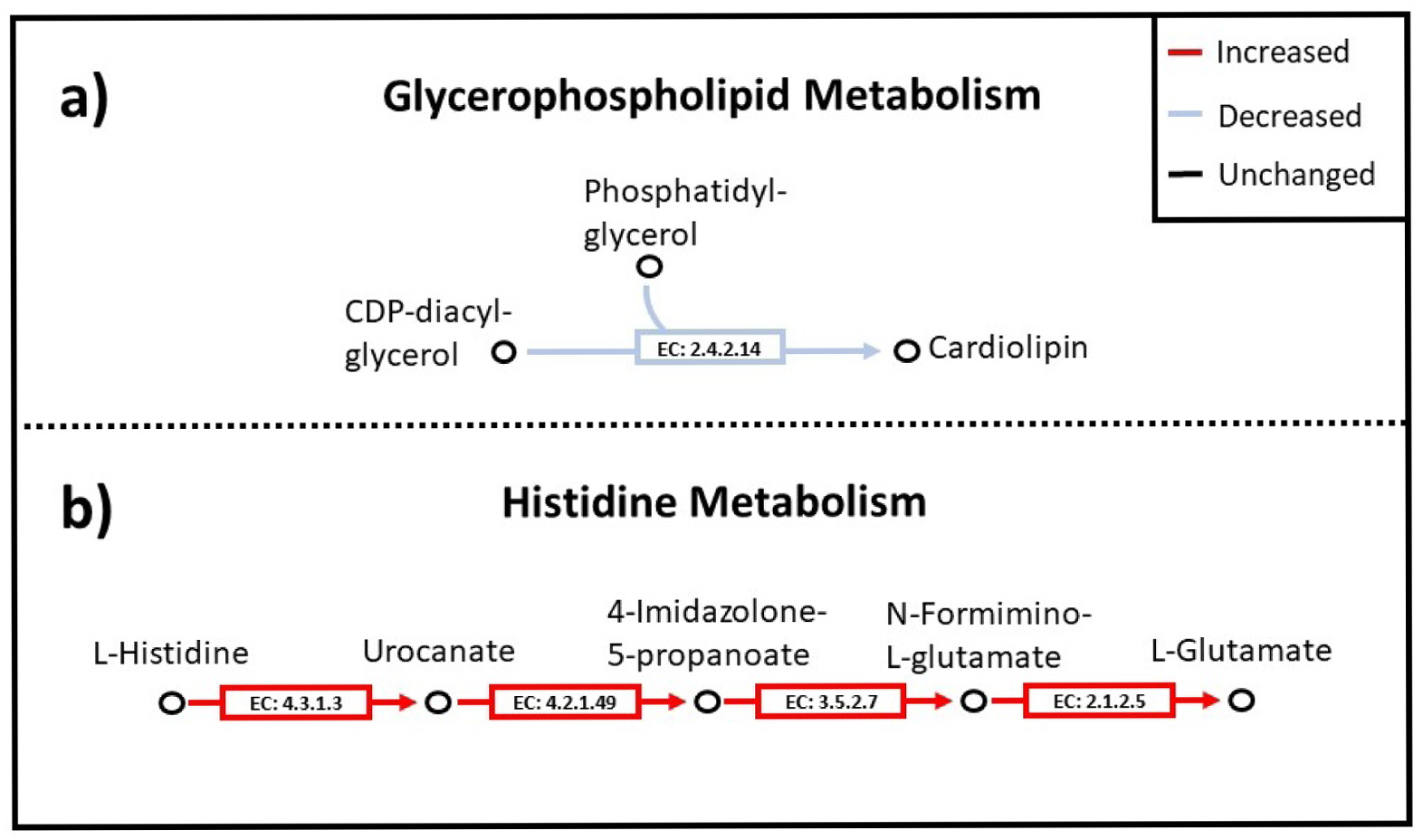
Reactions that show a notable flux change when the cell starts secreting D-serine. Reactions shown for: **a)** Glycerophospholipid metabolism, **b)** Histidine metabolism. The two cases compared were when D-serine secretion was 0 mmol/gDW hr. and 0.2 mmol/gDW hr.

Flux values for all the reactions in the model for these different cases from the FBA and the upper and lower bounds from the FVA analysis can be found in the supplementary material, along with some of the general unbound analysis cases.

### 3.3 Recombinant Protein Production

The results of the model optimisation for the five different recombinant proteins using the defined medium can be seen in Figure 7a. The different strain specific models all gave the same results as the general model. The case when the model was forced to keep at least 10% of the maximum biomass production gave less protein secretion for all types, since part of the resources were redirected from protein production and used to produce biomass. These can be found in the supplementary material. The case where D-serine was used to replace L-serine in the medium gave exactly the same results as the unchanged defined medium (Figure 7b). A combination of L- and D-serine was also tested, which again gave the same results, see supplementary material.

**Figure 7:**
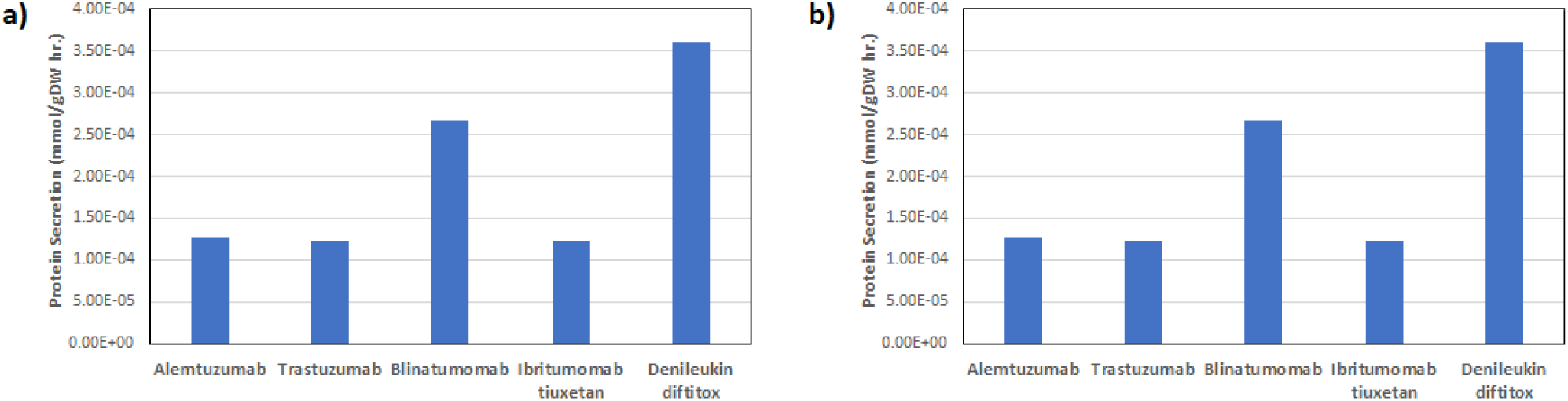
Optimisation of the production for the five different recombinant proteins using: **a)** defined medium and **b)** defined medium with D-serine instead of L-serine.

When analysing the reduced cost for the proteins, tryptophan appeared in the highest ten reactions for both ibritumomab tiuxetan and blinatumomab but not the other three. Reduced cost in this case simply refers to the derivative of the objective function with respect to the reactions, i.e. how much a change of a given reaction will affect the objective. This pointed to tryptophan amount, in the defined medium with L-serine, being a limiting factor for the production of these two proteins. To test this the tryptophan uptake rate in the defined medium was doubled. This led to increase in the secretion for both ibritumomab tiuxetan and blinatumomab, while the other proteins remained unchanged (Figure 8). Increasing the tryptophan amount even more did not result in any further changes.

**Figure 8:**
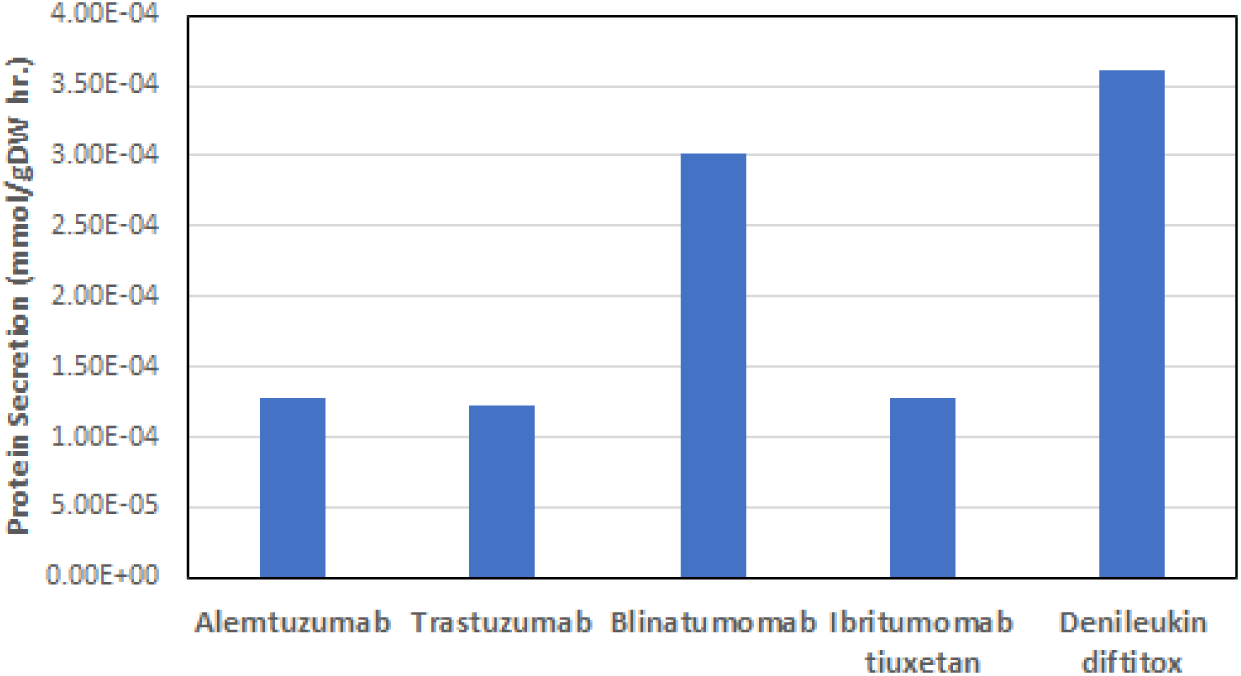
Optimisation of the production for the five different recombinant proteins using the defined medium with tryptophan intake doubled.

Repeating this type of procedure for blinatumomab revealed that increasing threonine and tyrosine uptake increased its production, while the increase for the other proteins was much more limited (Figure 9).

**Figure 9:**
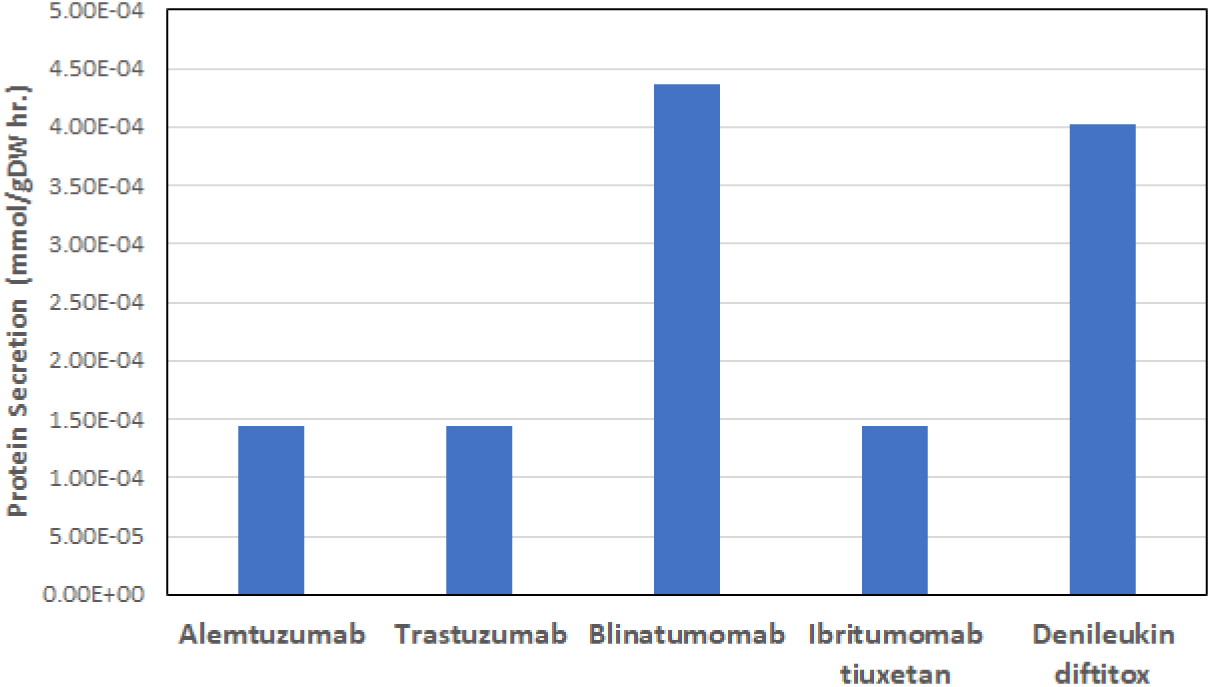
Optimisation of the production for the five different recombinant proteins using the defined medium with tryptophan, threonine and tyrosine intake doubled.

## 4 Discussion

### 4.1 Effect of NEAAs

The results in Figure 2 show that providing the cell with all the amino acids rather than only the essential ones enables considerably higher biomass production, even when the contents of the removed NEAAs are provided in the medium. This seems to show that it is more energy efficient for the cell to be provided with the NEAAs rather than having to utilise energy to generate them. Extracellular provision of NEAAs limits the extent of energy loss in amino acid biosynthetic pathways and renders additional reducing power available for other biological processes in the cell. Consequently, the amino acid biosynthetic routes do not impose additional burden on the CHO cell metabolism, allowing it to avoid potential constraints on the growth yield and cellular productivity.

The nine reported EAAs in the model are the same as the ones determined essential in humans. However, this is not in complete agreement with the literature. Along with these nine amino acids, a few others have been named as being essential in CHO cells. Arginine has for example been reported as being essential in CHO cells, and both cysteine and proline have been reported as CHO-specific auxotrophies [19]. These amino acids are not essential for growth using the general model, however, they are essential in the strain specific models. Adding these three to the EAA group and again removing the NEAAs, while replacing their contents in the medium, gave the exact same results as before, see supplementary material.

The results in Figure 3 show that only the removal of tyrosine and cysteine has any effect on the biomass production. For the other amino acids tested, it seems that the system can make up for a variation in one by automatically adjusting it using interconversion with other metabolites. These results are particularly interesting in light of the different biosynthetic cost of NEAAs. In both *E.coli* [26] and yeast [27] the two NEAAs with the highest biosynthetic costs are tyrosine and cysteine. However, this is not the case for human and animal cells where tyrosine is ranked considerably lower in regards to biosynthetic cost [27]. This is due to the fact that cysteine and tyrosine are synthesised from EAAs methionine and phenylalanine in these cases and cannot be synthesised *de novo* in animals. Since these precursors are EAAs that cannot be synthesised by the cells, its cost is not considered in the cost calculations, causing tyrosine to have much lower biosynthetic cost in mammalian cells. The other NEAAs can all be synthesised from basic metabolites produced during glycolysis and the Krebs cycle.

Forcing the cell to use these EAAs for this purpose is a major constraint on the metabolism. Although glucose and other such factors in the medium are increased to make up for the contents of the NEAAs that get taken out, the cell is still given the same limited amount of methionine and phenylalanine, forcing it to use that in NEAA synthesis rather than other purposes. For example, methionine is the universal initiator of protein synthesis and, therefore, extremely important in the metabolism. Analysis of extant proteomes has also shown that phenylalanine, cysteine and tyrosine have substantially increased in frequency within proteins since the three primary lineages diverged more than three billion years ago [28]. These results show that including NEAAs in the medium has a positive effect on the biomass production and that cysteine as well as tyrosine play a fundamental role in this.

### 4.2 D-serine Secretion

The D-serine secretion only appeared in extreme unbound cases, showing that this is likely not something that happens to the cell under natural medium conditions. The precise mechanism of D-serine release still remains unclear though and some reports suggest that it could be promoted by extracellular L-serine via transporter exchange or by sodium influx via sodium-dependent transporters [29]. It would, therefore, be interesting to see if such conditions might occur in any industrial or production circumstances.

To make sure the observed difference seen in Figures 5 and 6 between the non-secreting and secreting cases for D-serine was not merely because the amount of L-serine became limited, another test with extra L-serine input was performed. This test revealed that the 32 reactions still seemed to show a notable difference according to FVA, even though the biomass did not decrease with the increased L-serine available. However, during further analysis it was later discovered that the flux variability function in the COBRA package, that was used for the FVA calculations, was giving off wrong results when it was used on a subset of the reactions. This eventually led to the discovery of a bug in the function code that was subsequently fixed following the matter being brought up with the COBRA developers. More detailed information on this can be found in the supplementary material. When the test case was repeated it again showed that with increased L-serine the biomass production did not decrease, however, the 32 reactions subsequently did not show a notable difference anymore. Therefore, it seems that the observed flux differences only become significant when L-serine becomes the limiting factor when too much of it is forced into D-serine secretion, making the biomass production decrease. Thus, the metabolic flux changes can be traced to the system adjusting for this decrease in production. The decrease in production also explains the decrease in ATP consumption, since flow through purine and biomass production is reduced as L-serine becomes limiting.

However, the results in Figures 5 and 6 still show interesting correspondence to reported literature. For example, the reduced flux towards the sphingolipid metabolism is in agreement with reports that state that D-serine inhibits the pathway [30]. The increased flow from tryptophan could likewise be linked to reports of it being an inhibitor of D-serine [31]. Another study states that control of D-serine secretion is directly related to activation and deactivation of certain steps within glycolysis and the Krebs cycle, however, no flux changes observed here seemed to show such indication [32].

The increased flux from histidine to glutamate is also interesting since D-serine is an essential co-agonist with glutamate for the stimulation of N-methyl D-aspartate (NMDA) receptors in the brain [33]. D-serine has been the object of several studies to ascertain the nature of its metabolism as it is a crucial signalling molecule utilised by astroglia and neurons in the mammalian central nervous system. It is of high interest since it has been linked to various medical conditions, for example decreased D-serine levels have been associated with the hyperfunction of NMDA receptors in schizophrenia as well as the cognitive impairments observed during aging [33]. Elevated D-serine levels in the spinal cord of patients with amyotrophic lateral sclerosis are also believed to mediate motor neuron degeneration [33, 34]. Metabolic modelling approaches such as used in this study could, therefore, perhaps be a valuable tool to help gain deeper understanding of its metabolism in these cases.

### 4.3 Recombinant Protein Production

The results of the model optimisation for the different recombinant proteins seen in Figure 7a show clearly how the difference in protein structure can influence the production outcome. The secretion varies greatly depending on the protein, with the highest secretion being for denileukin diftitox and lowest for trastuzumab. The fact that the outcome does not change depending on which strain was used shows that the medium rather than strain seems to have the limiting effect in this case, although of course not all these recombinant proteins would be well suited for production in all the different strains. The size of the proteins is obviously a big factor here, with the smaller proteins, denileukin diftitox and blinatumomab, having much larger flux values than the three other larger proteins. However, size alone does not explain the secretion difference since denileukin diftitox has considerably higher production than blinatumomab, despite being larger. It is, therefore, clear that the amino acid content also has a large effect. For the larger three proteins the difference is not as obvious, however, ibritumomab tiuxetan secretion is almost the same as for trastuzumab, even though ibritumomab tiuxetan is smaller. Looking at the amino acid compositions of the different proteins it can be seen that the tryptophan proportion of both ibritumomab tiuxetan and blinatumomab are relatively higher than for the other proteins (Table 3).

**Table 3:** Tryptophan content for each recombinant protein

|  | **Tryptophan content**<br>**of protein** |
| --- | --- |
| **Alemtuzumab** | 1.51% |
| **Trastuzumab** | 1.66% |
| **Blinatumomab** | 2.58% |
| **Ibritumomab**<br>**Tiuxetan** | 2.15% |
| **Denileukin**<br>**Diftitox** | 0.96% |

The results in Figure 8 indicate that in the defined medium, tryptophan becomes a limiting factor for the production of ibritumomab tiuxetan and blinatumomab. This is, therefore, likely caused by the relatively high tryptophan proportion of both proteins. The results in Figure 9, indicate that threonine and tyrosine are limiting factors for the production of blinatumomab. However, what is interesting is that threonine composition in blinatumomab is not particularly high compared to the other proteins, emphasising that blindly increasing the amino acids with the highest composition in the target protein is not necessarily the best tactic. Optimising the amino acid media composition is a multi-parametric problem which is why taking metabolic complexity into account can be of great help narrowing down the search space for experimental analysis.

The fact that the results remain the same for the protein production when the L-serine is swapped for D-serine in the defined medium as seen in Figure 7b is quite interesting. This seems to be in agreement with a hypothesis put forward by Puyal *et al.* that neurons can use D-serine metabolism as a supplementary energetic pathway [35]. This would be interesting to confirm with experiments.

In summary, the results presented here show how the system can be utilised in broad *de novo* exploration of amino acid content as well as tailoring medium composition for protein production depending on the protein’s structure and size. As mentioned, the system could also be relevant as a possible model system to gain deeper understanding of more complex mammalian cells. However, given the variability of the model as seen in the FVA analysis carried out in this study, the model would have to be constrained considerably to get a better large-scale picture of notable changes in the cell metabolism. As access to available omics data continues to grow this will become easier. This model variability also shows the cellular robustness of the system and how the cell has great flexibility in its metabolism to react to different environments and salvage itself. This is for example something that small-scale kinetic models would not pick up on.

A possible set-back of the analysis presented in this study is that the used biomass objective function was defined using previous experimental measurements from the literature [19]. However, the composition of biomass from mammalian cells is not static and, therefore, this might impact model predictions of growth and recombinant protein production. This problem could be improved by carrying out more comprehensive measurements of the composition of CHO cells under various conditions and use it to define a more accurate biomass objective function.

## 5 Conclusion

Exploratory analysis of the amino acid requirements of CHO cells was carried out using metabolic modelling approaches. The results showed that including NEAAs in the medium has a positive effect on the biomass production and that cysteine as well as tyrosine play a fundamental role in this. The reason for this seems to stem from the fact that extracellular provision of NEAAs limits the extent of energy loss in amino acid biosynthetic pathways and renders additional reducing power available for other biological processes. Detailed analysis of the possible secretion and uptake of D-serine in the CHO model was also performed and its influence on the rest of the metabolism mapped out. This revealed some results matching various existing literature, as well as leading to the discovery of a bug in the Cobra toolbox. These results are interesting since no mention of D-serine in regard to CHO cells was found in current literature, as well as the fact that this opens up the possibility of using the model for higher up organisms. Finally, results for the model optimisation of different recombinant proteins showed clearly how the difference in protein structure and size can influence the production outcome. Limiting factors in the defined medium were identified for both ibritumomab tieuxetan and blinatumomab as well as it was shown how their increase led to improved protein secretion. In summary, the results show that systematic and model-based approaches such as the ones used in this study indicate great possibility for broad *de novo* exploration as well as being able to handle the cellular burden associated with the production of different types of recombinant protein. Implementing these types of approaches as a long-term investment could, therefore, be of great advantage for process development of biopharmaceuticals.

## 6 Acknowledgements

I gratefully acknowledge support from the Bill & Melinda Gates Foundations through the Gates-Cambridge Scholarship under the grant number OPP1144.

## 7 Supplementary Material

Supplementary material can be accessed at the following online directory: https://doi.org/10.17863/CAM.44558

